# Tonic GABAergic activity facilitates dendritic calcium signaling and short-term plasticity

**DOI:** 10.1101/2020.04.22.055137

**Authors:** Chiayu Q. Chiu, Thomas M. Morse, Francesca Nani, Frederic Knoflach, Maria-Clemencia Hernandez, Monika Jadi, Michael J. Higley

## Abstract

Brain activity is highly regulated by GABAergic activity, which acts via GABA_A_Rs to suppress somatic spike generation as well as dendritic synaptic integration and calcium signaling. Tonic GABAergic conductances mediated by distinct receptor subtypes can also inhibit neuronal excitability and spike output, though the consequences for dendritic calcium signaling are unclear. Here, we use 2-photon calcium imaging in cortical pyramidal neurons and computational modeling to show that low affinity GABA_A_Rs containing an α5 subunit mediate a tonic hyperpolarization of the dendritic membrane potential, resulting in deinactivation of voltage-gated calcium channels and a paradoxical boosting of action potential-evoked calcium influx. We also find that GABAergic enhancement of calcium signaling modulates short-term synaptic plasticity, augmenting depolarization-induced suppression of inhibition. These results demonstrate a novel role for GABA in the control of dendritic activity and suggest a mechanism for differential modulation of electrical and biochemical signaling.

## Introduction

Neuronal activity in the mammalian brain is strongly shaped by GABAergic signaling, mediated by the release of GABA from local interneurons and the subsequent binding to both ionotropic type-A receptors (GABA_A_Rs) and metabotropic type-B receptors (Farrant and Nusser, 2005). Synaptic GABAergic inhibition through both receptor types regulates the timing and magnitude of action potential generation and also shapes synaptic integration and calcium signaling in dendrites (Cardin, 2018; Higley, 2014). Indeed, we and others previously showed that synaptic activation of GABA_A_Rs suppressed calcium influx through both voltage-gated calcium channels and NMDA-type glutamate receptors (Chiu et al., 2013; Marlin and Carter, 2014). Moreover, GABAergic control of dendritic calcium is a critical modulator of synaptic plasticity (Cichon and Gan, 2015; Hayama et al., 2013). Finally, disruption of GABAergic activity is implicated in a variety of neuropsychiatric disorders, where an imbalance of synaptic excitation and inhibition is argued to drive network dysfunction (Gogolla et al., 2009).

In addition to synaptic transmission, GABA can also evoke a tonic membrane conductance thought to be mediated in part via extrasynaptic GABA_A_Rs with distinct subunit composition than their synaptic counterparts (Bai et al., 2001; Farrant and Nusser, 2005; Scimemi et al., 2005; Stell and Mody, 2002). In the hippocampus and neocortex, α4,5,6 and δ subunits have been most closely linked to tonic GABAergic conductances (Jacob, 2019; Lee and Maguire, 2014). Tonic current through GABA_A_Rs can suppress neuronal output by hyperpolarizing the membrane potential and shunting the integration of synaptic inputs (Farrant and Nusser, 2005; Jacob, 2019; Lee and Maguire, 2014). However, the consequences of tonic GABA_A_R activity for dendritic calcium signaling are unclear.

Here, we used 2-photon laser scanning microscopy (2PLSM) to investigate the role of GABAergic signaling in apical dendrites of layer 2/3 neurons from the mouse prefrontal cortex. We found the surprising result that blocking GABA_A_Rs paradoxically suppressed dendritic calcium influx in response to somatic action potentials (APs). This effect was mimicked by selective decrease in the activity of low-affinity, α5 subunit-containing receptors. In contrast, application of a novel α5-specific positive modulator enhanced dendritic calcium influx. GABAergic regulation of dendritic calcium signaling also modulated the short-term plasticity of synaptic inhibition. Computational modeling revealed that tonic GABAergic hyperpolarization of the dendritic membrane potential deinactivates low-threshold calcium channels and boosts their subsequent activation by back-propagating APs, a hypothesis confirmed experimentally. Overall, these counterintuitive results demonstrate that tonic GABAergic signaling can enhance dendritic calcium influx and short-term synaptic plasticity. Our work highlights a novel role for α5 subunit-containing receptors in the cortex and suggests new avenues for the exploration of GABAergic control of neuronal activity in both health and disease.

## Results

In order to investigate the role of GABAergic signaling on dendritic activity, we performed whole-cell recordings from layer 2/3 pyramidal neurons (PN_S_) in acute slices of mouse medial prefrontal cortex. Cells were filled through the patch pipette with the structural indicator Alexa Fluor 594 and the calcium indicator Fluo 5F and imaged using 2PLSM (Figure 1A). Somatic APs that propagated into the dendritic arbor were evoked using brief current injection. The resulting calcium transients were visualized in single dendritic spines and neighboring dendritic shafts in line-scan mode (Figure 1A-B).

**Figure 1.**
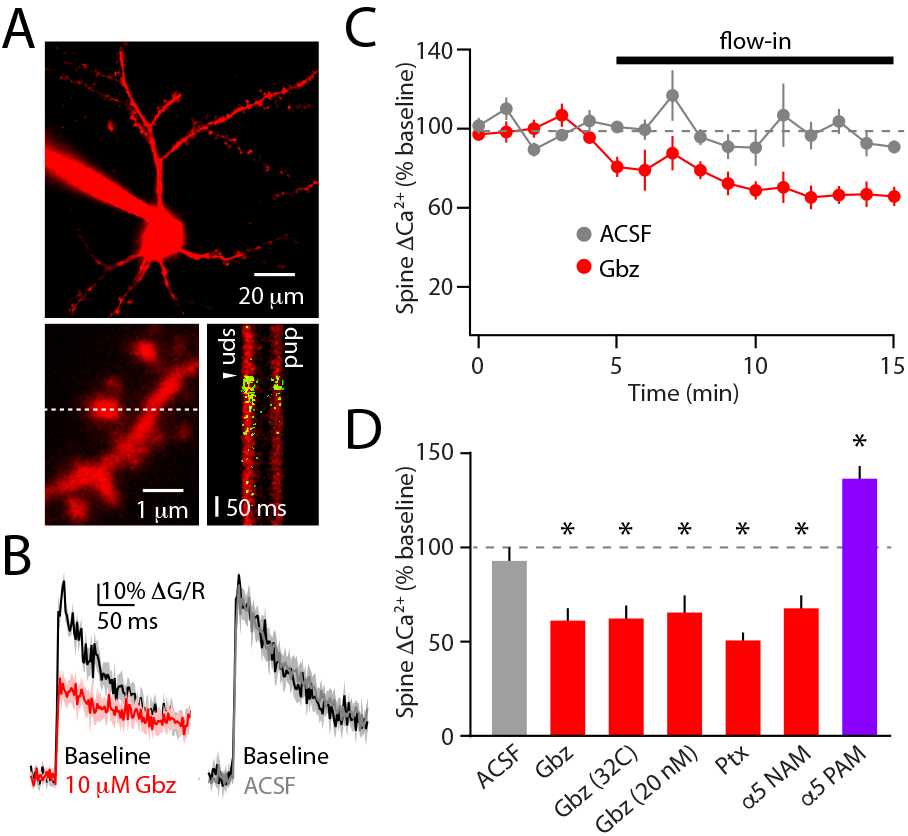
GABAergic signaling enhances action potential-evoked dendritic calcium transients. **A**, Representative image showing the recording configuration. Layer 2/3 PNs from the mouse prefrontal cortex were filled through the patch pipette with Alexa Fluor 594 and Fluo 5F and imaged using 2-photon laser-scanning microscopy (top). Somatic action potentials were evoked via somatic current injection (time indicated by arrowhead), producing brief dendritic calcium transients in spines and neighboring shafts (dashed line indicates line-scan), visible as a rise in green fluorescent signal (bottom). **B**, Average AP-evoked calcium transients measured in dendritic spines under baseline conditions (Ctl, black) and after flow-in of 10 μM gabazine (red, left traces) or ACSF (gray, right traces). Lines and shaded regions indicate mean ± SEM. **C**, Average time-course of reduction in AP-evoked spine ΔCa^2+^ relative to baseline (dashed line) following flow-in of 10 μM gabazine (red) or ACSF (gray), corresponding to data in (B). Error bars indicate SEM. **D**, Population data indicating the change in spine ΔCa^2+^ following flow-in of ACSF, 10 μM gabazine, 10 μM gabazine at 32°C, 20 nM gabazine, 10 μM picrotoxin, 10 μM RO4938581 (NAM), or 1 μM Compound A (PAM). Error bars indicate SEM. * indicates p<0.05 relative to baseline (dashed line).

To block GABAergic signaling, we bath-applied the GABA_A_R antagonist gabazine and observed a substantial reduction in the magnitude of the AP-evoked calcium transient (ΔCa^2+^). This reduction in ΔCa^2+^ was not caused by rundown, as control experiments applying artificial cerebrospinal fluid (ACSF) did not result in a significant change in transient amplitude (Figure 1B-C). Relative to pre-drug baseline, 10 μM gabazine (61.5±6.0%, n=13 spines, Wilcoxon Test, p=0.0002) but not ACSF (97.7±4.8%, n=5 spines, Wilcoxon Test, p=0.62) produced a significant reduction in ΔCa^2+^. We observed similar results when experiments were carried out at near-physiological temperature (62.2±6.2%, n=8 spines, Wilcoxon Text, p=0.008) and with the distinct GABA_A_R blocker picrotoxin (50.9±3.8%, n=13 spines, Wilcoxon Test, p=0.0002, Figure 1D). Moreover, ΔCa^2+^ was also reduced when measured in the dendritic shaft adjacent to each spine (Supplemental Figure 1). Alterations in calcium signals occurred with minimal changes to either the resting membrane potential or input resistance wen measured from the somatic recording site (Supplemental Figure 1).

In the neocortex, GABA_A_Rs containing α5 subunits have been linked to tonic currents and exhibit a relatively lower affinity for GABA (Farrant and Nusser, 2005; Jacob, 2019; Scimemi et al., 2005), suggesting they may be sensitive to reduced concentrations of competitive antagonists. Consistent with this hypothesis, we found that application of 20 nM gabazine produced a similar reduction in ΔCa^2+^ (65.9±9.0%, N=8 spines, Wilcoxon Test, p=0.016, Figure 1D). We next tested the α5 subunit-specific negative allosteric modulator (NAM) RO4938581 (Ballard et al., 2009) and again observed a significant reduction in ΔCa^2+^ (66.8±7.9%, n=7 spines, Wilcoxon Test, p-0.006, Figure 1D). Finally, we reasoned that if suppression of α5 subunit-containing receptors could reduce dendritic calcium signals, enhancement of their activity might have the opposite effect. Therefore, we took advantage of a novel positive allosteric modulator (PAM) developed by Roche termed Compound A, which exhibits a high degree of specificity for α5 subunit-containing GABA_A_Rs (Supplemental Figure 2). Consistent with our hypothesis, application of Compound A produced a significant increase in ΔCa^2+^ (135.5±8.3%, n=7 spines, Wilcoxon Test, p=0.005, Figure 1D). As above, α5-specific manipulations did not alter somatic membrane potential or input resistance (Supplemental Figure 1).

Dendritic calcium signaling evoked by depolarization is linked to various forms of synaptic plasticity, including depolarization-induced suppression of inhibition (DSI). DSI occurs when postsynaptic spiking drives calcium-dependent dendritic endocannabinoid synthesis and subsequent retrograde suppression of GABA release from presynaptic terminals (Chevaleyre et al., 2006). We established a DSI induction protocol (50 APs at 50 Hz) using brief current pulses through the somatic pipette and observed a robust dendritic calcium signal that was significantly reduced by application of 20 nM gabazine (71.6±7.7% relative to pre-drug baseline, n=6 spines, Wilcoxon Signed Rank Test, p=0.03, Figure 2A-B). We evoked inhibitory postsynaptic currents (IPSCs) using local electrical stimulation and found that DSI induction produced a significant, transient suppression of IPSC amplitude relative to the pre-induction period (80.5±2.9%, n=8 cells, Wilcoxon Signed Rank Test, p=0.008, Figure 2C-D). Application of 20 nM gabazine reduced but did not block evoked IPSC amplitude (52.8±6.4% relative to pre-drug baseline, Figure 2C). However, gabazine significantly decreased the amount of DSI observed (90.7±2.2%, Wilcoxon Signed Rank Test relative to pre-induction, p=0.02; Wilcoxon Matched-Pairs DSI in gabazine versus pre-drug baseline, p=0.02, Figure 2D-E). These data suggest that the reduction in dendritic calcium signaling associated with loss of tonic GABAergic activity results in weaker induction of calcium-dependent short-term plasticity.

**Figure 2.**
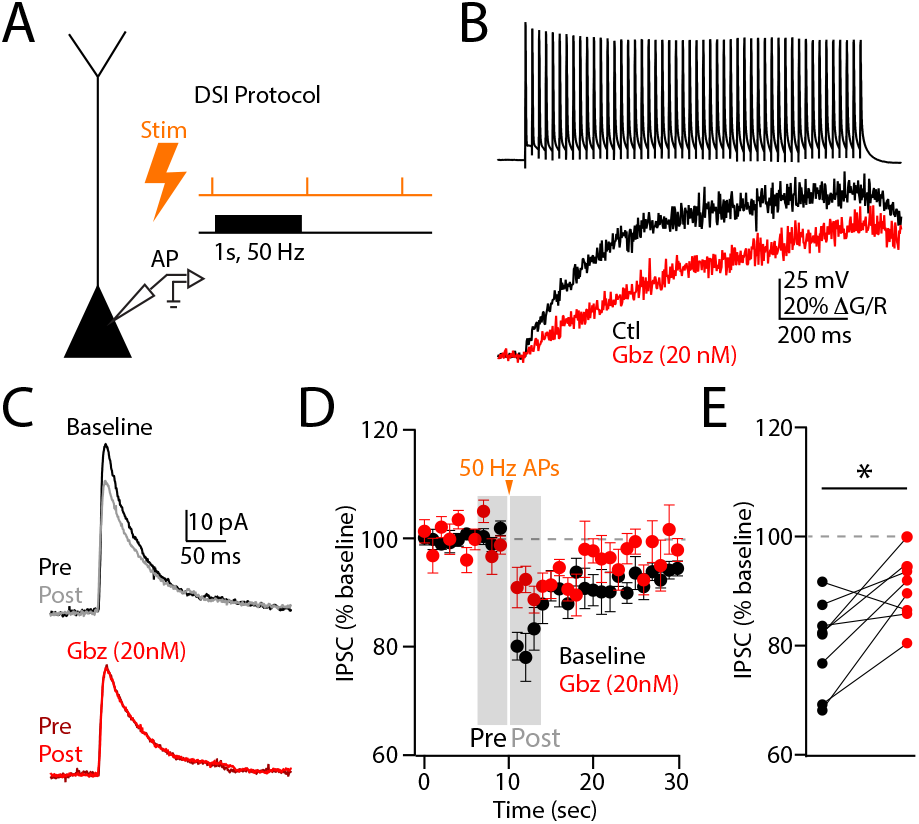
Tonic GABAergic signaling enhances depolarization-induced suppression of inhibition. **A**, Schematic illustration of the experimental setup. Whole-cell recordings were obtained from layer 2/3 PNs. Local electrical stimulation was used to evoke IPSCs. DSI was induced with a brief train of somatic APs (1 second, 50 Hz). **B**, Low concentration (20 nM) gabazine reduces ΔCa^2+^ in dendritic spines evoked by train of somatic APs used for DSI induction. **C**, Example IPSCs recorded before (Pre) or after (Post) DSI induction, either under baseline conditions (black) or after flowin of 20 nM gabazine (red). **D**, Average time-course of IPSC amplitude relative to baseline (dashed line) before and after DSI induction, recorded either under baseline conditions (black) or after flow-in of 20 nM gabazine (red). DSI induction time is indicated by arrowhead. Time windows for calculating Pre and Post IPSC amplitudes are shown in gray. Error bars indicate SEM. **E**, Population data showing the average DSI (measured as % baseline IPSC amplitude) before (black) or after (red) gabazine flow-in. * indicates p<0.05 relative to baseline (dashed line).

We next explored the biophysical mechanisms that might mediate the enhancement of dendritic calcium signaling via tonic GABAergic activity. We simulated a biophysically realistic neuron that included a cell body and dendritic arbor (Supplemental Figure 3, see Methods). To accurately reflect the expression of diverse voltage-gated calcium channels in cortical neurons, we included both high (“HVA”, N/P/Q-like) and low (“LVA”, T-like) conductances. These channel types differ primarily in the dependence of their activation and inactivation dynamics on membrane voltage (Supplemental Figure 3). Finally, we simulated a tonic GABAergic conductance uniformly distributed in the cell membrane with a reversal potential of −70 mV, matching our previous data from perforated patch recordings in these cells (Chiu et al., 2013).

Under control conditions (GABA conductance intact), simulating an evoked somatic action potential strongly depolarized the dendritic compartment, producing a calcium current through both HVA and LVA channels (Figure 3A). We then repeated the simulation with the GABA conductance blocked, mimicking our experimental gabazine conditions. As with the experimental data, loss of GABAergic activity led to a ~35% reduction in peak calcium current. Examination of the different calcium channel subtypes revealed a modest (~15%) reduction in peak HVA conductance and a much larger (~68%) reduction in peak LVA conductance. The decreased LVA conductance was primarily mediated by an increased resting channel inactivation associated with depolarization of the dendritic membrane potential following GABA_A_R blockade (Figure 3A). Notably, the model confirmed that the somatic membrane potential and input resistance were only minimally altered, consistent with our experimental data (Figure 3B). Further examination of the relationship between membrane potential and dendritic location revealed that the tonic GABA conductance produced an increasing hyperpolarization with distance from the cell body relative to the blocked condition, contributing to deinactivation of LVA calcium channels in the more distal dendrites (Figure 3C). Consistent with this conclusion, the reduction in evoked calcium transient following GABA blockade was enhanced for more distal dendritic locations and was greater when the model incorporated exclusively LVA versus HVA conductance (Figure 3D).

**Figure 3.**
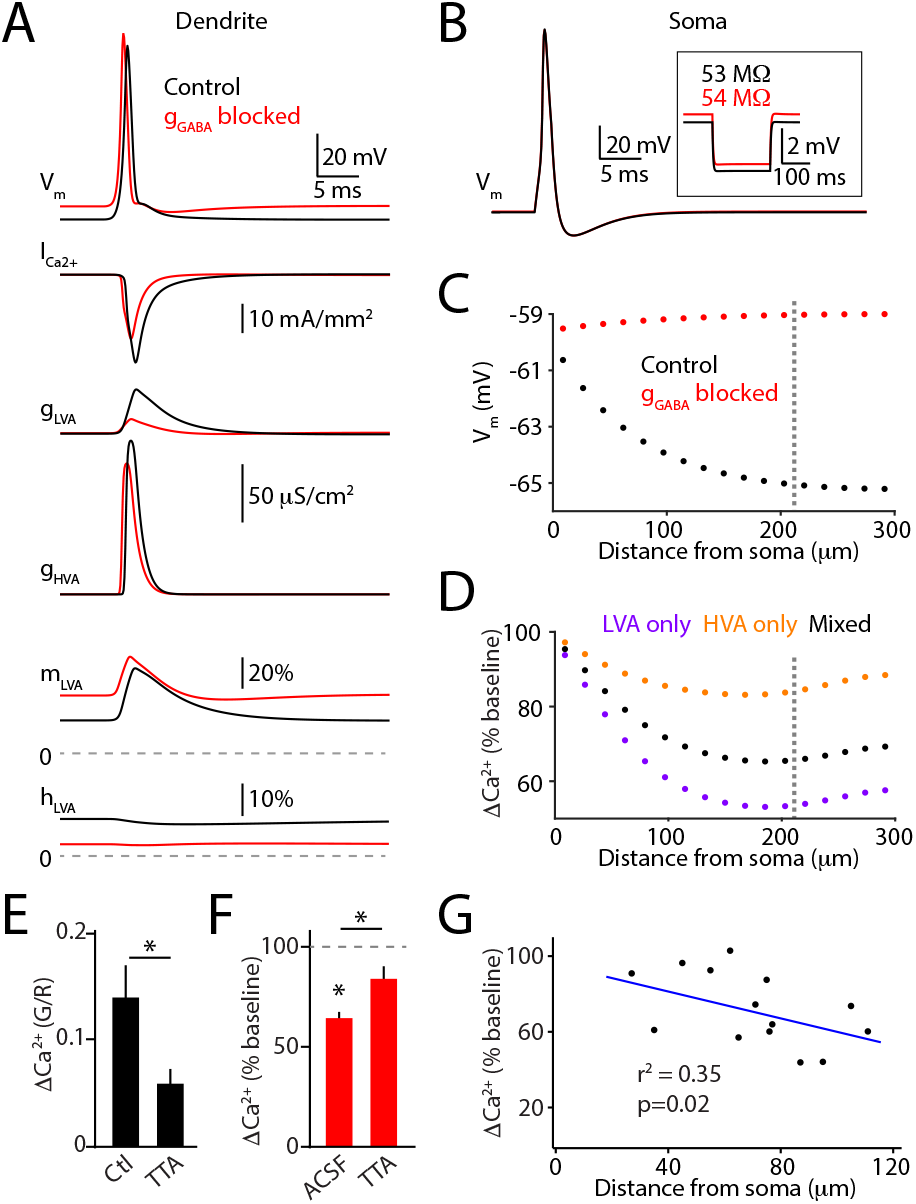
Tonic GABAergic signaling deinactivates dendritic low-threshold voltage-gated calcium channels. **A**, Computational model data illustrating dendritic voltage (V_m_), calcium current (I_ca2+_), low- and high-threshold calcium channel conductance (g_LVA_, g_HVA_), and low-threshold activation and inactivation gate open fraction (m_LVA_, h_LVA_) in response to a somatic action potential. Data are shown for control conditions (black) and following blockade of tonic GABAergic conductance (red). **B**, Computational model data illustrating somatic voltage (V_m_) in response to evoked action potential under control conditions (black) and following blockade of tonic GABAergic conductance (red). Inset shows somatic membrane potential response to brief hyperpolarizing current pulse in the two conditions and the corresponding input resistance values. **C**, Computational data showing membrane voltage (V_m_) as a function of distance from the soma under control conditions (black) and following blockade of GABAergic conductance (red). Dendritic location for data in (A) is shown by vertical dashed line. **D**, Computational data showing change in ΔCa^2+^ relative to baseline following blockade of GABAergic conductance. Data are shown for model versions with low-threshold (LVA only, purple), high-threshold (HVA only, orange), or combination of channels (Mixed, black). Data from (A) were acquired with the Mixed version at the indicated dendritic location (vertical dashed line). **E**, Experimental data showing application of the selective T-type calcium channel blocker TTA-A2 (TTA) significantly reduces AP-evoked ΔCa^2+^ in dendritic spines. **F**, Experimental data showing that blockade of T-type calcium channels by pre-incubation with TTA-A2 (TTA) significantly eliminates the reduction in ΔCa^2+^ relative to baseline (dashed line) caused by flow-in of 10 μM gabazine. **G**, Experimental data showing that the magnitude of ΔCa^2+^ relative to baseline caused by gabazine flow-in is significantly correlated with increasing distance from the soma (blue line indicates linear fit).

These computational findings suggest that the low-threshold inactivation of the LVA channels is the primary mechanism underlying the interaction of GABAergic signaling and dendritic calcium influx and that the reduction in ΔCa^2+^ following application of GABAergic antagonists should be greater for more distal dendritic locations. We therefore analyzed dendritic activity in the presence of the selective T-type channel blocker TTA-A2. Application of TTA-A2 alone significantly reduced ΔCa^2+^ in dendritic spines (Control ΔG/R: 0.14±0.03 n=9 spines, TTA-A2 ΔG/R: 0.06±0.01 n=6 spines, Mann-Whitney Test, p=0.036, Figure 3E), consistent with previous reports that these channels make a substantial contribution to AP-evoked dendritic calcium influx in the prefrontal cortex (Chalifoux and Carter, 2011). Furthermore, in contrast to the effect in control cells (65.0±4.1%, n=9 spines, Wilcoxon Test, p=0.004), pre-incubation of cells in TTA-A2 eliminated the significant reduction in calcium signal associated with application of gabazine (83.8±7.7%, n=6 spines, Wilcoxon Test, p=0.16; Mann-Whitney Test comparing ACSF with TTA-A2, p=0.026, Figure 3F). Finally, we examined the relationship between the gabazine-induced reduction in calcium signal and dendritic location, finding a significant correlation between these values (Spearman’s rank correlation, r=-0.59, p=0.02, Figure 3G), with suppression increasing in more distal regions. Thus, the experimental data provide direct confirmation of mechanisms suggested by our computational model, indicating that a tonic GABAergic conductance can hyperpolarize the distal dendritic membrane potential, deinactivating low-threshold calcium channels and enhancing AP-evoked calcium influx.

## Discussion

In the present study, we show that GABAergic hyperpolarization of cortical dendrites surprisingly enhances AP-evoked calcium signals as well as calcium-dependent short-term plasticity of synaptic inhibition. Pharmacological evidence suggests this result is mediated by the activity of α5 subunit-containing GABA_A_Rs, and dendritic calcium influx can be bi-directionally controlled by α5-selective negative and positive modulators. Our computational modeling and experimental data indicate a critical role for low-threshold calcium channels (e.g., T-type), with greater GABAergic modulation occurring with increasing distance from the cell body. Strikingly, we also found that tonic GABAergic boosting of dendritic calcium influx can enhance DSI, a calcium-dependent form of short-term inhibitory synaptic plasticity (Chevaleyre et al., 2006). These results support the conclusion that dendritic and somatic compartments are not isopotential and further demonstrate the complexity of signal integration along the somatodendritic axis.

Previous studies have clearly demonstrated a role for tonic GABAergic conductances in the steady suppression of neuronal output, mediated by both membrane hyperpolarization and shunting of synaptic inputs (Farrant and Nusser, 2005; Lee and Maguire, 2014). Indeed, α5 subunit-containing GABA_A_Rs were recently shown to suppress the back-propagation of APs in PN dendrites and the opening of NMDA-type glutamate receptors (Groen et al., 2014; Schulz et al., 2018). Our present results are not inconsistent with this view. However, the relationship between membrane potential and channel inactivation adds additional complexity to the consequences of GABAergic signaling. For example, suppression of spiking (or dendritic propagation) may co-exist with a boosting of calcium influx driven by a greater fraction of channels in the deinactivated state. The exact nature of this complex relationship will certainly depend on a variety of factors, including instantaneous membrane potential, GABA_A_R reversal potential, and the specific voltage-dependent dynamics of dendritic calcium channels. As shown with our data, these factors are likely to vary along the dendritic arbor, providing a substrate for highly localized synaptic and extrasynaptic functions for GABAergic signaling (Higley, 2014).

The role of α5 subunit-containing receptors in our study adds to a growing literature on the functional roles of these channels in both health and disease (Ballard et al., 2009; Engin et al., 2018; Hauser et al., 2005; Paine et al., 2020). Previous studies have shown the involvement of α5 subunits in tonic inhibitory currents recorded in the hippocampus and neocortex (Caraiscos et al., 2004; Jacob, 2019). Pharmacological or genetic manipulation of α5 activity can produce significant modulation of learning and cognitive ability (Braudeau et al., 2011; Collinson et al., 2002; Donegan et al., 2019; Magnin et al., 2019). Altered α5 expression has also been linked to neuropsychiatric disorders, and α5 modulators may have antidepressant actions (Carreno et al., 2020; Prevot et al., 2019). Surprisingly, both negative and positive modulators of α5 subunit-containing receptors have been linked to therapeutic and pro-cognitive effects. One potential resolution for these seemingly disparate findings is that suppression of spiking activity and boosting of dendritic calcium signaling may play distinct roles in modifying behavior. Our results suggest that adjunct targeting of voltage-gated calcium channels may provide a novel approach to enhance the behavioral consequences of α5-dependent behavioral modulation.

The impact of tonic GABAergic signaling on neuronal function may also be dynamic. Activation of NMDA-type glutamate receptors is implicated in plasticity of both synaptic and extrasynaptic GABAergic synapses (Chiu et al., 2019; Chiu et al., 2018; Gu et al., 2016). In addition, activation of kainate-type glutamate and muscarinic cholinergic receptors can also potentiate tonic inhibition in the hippocampus (Dominguez et al., 2016; Jiang et al., 2015). The interaction of these processes with GABAergic control of dendritic calcium signaling is unknown.

A critical open question is the membrane distribution of receptors mediating the tonic current driving our results. Receptors containing α5 subunits have been linked to both synaptic and extrasynaptic locations, and it is unclear how these potentially distinct populations contribute to tonic signals (Fritschy and Mohler, 1995; Magnin et al., 2019; Schulz et al., 2018). In addition, the sources of GABA that promote tonic signaling are unclear. Both somatostatin- and vasoactive intestinal peptide-expressing interneurons form synapses utilizing α5 subunits (Magnin et al., 2019; Schulz et al., 2018), suggesting they may also supply GABA for tonic activity.

In conclusion, our data indicate a novel role for GABAergic activity in the control of dendritic calcium influx and calcium-dependent plasticity. Our experimental and computational data demonstrate the complexity of interactions between electrical and biochemical signaling and suggest new possibilities for exploring GABAergic approaches to behavioral modulation in both healthy individuals and patients suffering from neurodevelopmental and neuropsychiatric disease.

## Methods

### Slice Preparation

All animal handling was performed according to the Yale Institutional Animal Care and Use Committee and federal guidelines. Acute slices of the prefrontal cortex were prepared from wild-type C57/Bl6 mice at postnatal day 25-40. Briefly, isoflurane-anesthetized mice were decapitated and 300 μm coronal slices were cut in ice-cold external solution containing (in mM): 100 choline, 25 NaHCO_3_, 1.25 NaH_2_PO_4_, 2.5 KCl, 7 MgCl_2_, 0.5 CaCl_2_, 15 glucose, 11.6 sodium ascorbate and 3.1 sodium pyruvate, bubbled with 95% O_2_ and 5% CO_2_. Slices containing prelimbic and infralimbic regions were then transferred to artificial cerebrospinal fluid (ACSF) containing (in mM): 127 NaCl, 25 NaHCO_3_, 1.25 NaH2PO4, 2.5 KCl, 1 MgCl2, 2 CaCl2 and 15 glucose, bubbled with 95% O2 and 5% CO2. After an incubation period of 30 min at 34 °C, the slices were maintained at 22–24 °C for at least 20 min before use.

### Electrophysiology and imaging

Experiments were conducted at room temperature (~22°C) or near-physiological temperature (32°C) where noted in a submersion-type recording chamber. Whole-cell recordings were obtained from layer 2/3 pyramidal cells (200-300 μm from the pial surface) identified with video-infrared/differential interference contrast. Glass electrodes (3.2-3.8 MΩ) were filled with internal solution containing (in mM): 135 KMeSO_3_, 10 HEPES, 4 MgCl_2_, 4 Na_2_ATP, 0.4 NaGTP and 10 sodium creatine phosphate, adjusted to pH 7.3 with KOH. Red-fluorescent Alexa Fluor-594 (10 μM) and green-fluorescent Fluo-5F (300 μM) were included in the pipette solution to visualize cell morphology and changes of intracellular calcium concentration, respectively. For non-imaging experiments (Figure 2), Fluo-5F was replaced with 100 μM EGTA. The series resistance was <25Ω and uncompensated. Recordings were discarded if the series resistance changed more than 20% during the experiment. Neurons were filled via the recording pipette for at least 15 minutes before imaging. To standardize recordings across cells, the membrane potential was adjusted to −60 mV by injection of small amounts of depolarizing current through the recording pipette. Electrophysiological recordings were made using a Multiclamp 700B amplifier, filtered at 4 kHz and digitized at 10 kHz.

Imaging was performed with a custom-built microscope, including components manufactured by Mike’s Machine Company. Fluorophores were excited using 840 nm light from a titanium-sapphire laser (Ultra-2, Coherent). Emitted green and red photons were separated and collected by photomultiplier tubes. Imaged spines were located along secondary and tertiary branches of the apical dendrite. Action potentials were evoked using a brief depolarizing current pulse (1 ms, 2 nA) through the recording pipette. For imaging AP-evoked transients, signals were collected during 500 Hz line scans across a spine and the neighboring dendritic shaft. Reference frame scans were taken between each acquisition to correct for small spatial drift of the preparation over time. Calcium signals were quantified as increases in green fluorescence from baseline normalized to the average red fluorescence (ΔG/R).

For DSI experiments, we placed a glass theta stimulating electrode in layer 1 < 100 μm from the recorded cell and used brief (0.1 ms) pulses to evoke IPSCs every 6 s. DSI was induced with action potential bursts evoked by somatic current injection (as above) delivered at 50 Hz for 1 s in the presence of a low concentration of the metabotropic glutamate receptor agonist DHPG.

### Data acquisition and analysis

Data were acquired using National Instruments data acquisition boards and ScanImage software (Pologruto et al., 2003). Off-line analysis was performed using custom routines written in MATLAB (The Mathworks) and IgorPro (Wavemetrics). AP-evoked ΔCa^2+^ was calculated as the average ΔG/R over a 50 ms window, starting 5 ms after the stimulus. IPSC amplitudes were calculated by finding the peak of the current traces and averaging the values within a 1 ms window. To assess DSI, we averaged IPSCs 4 trials before and after induction.

### Pharmacology

For all experiments, except where noted, the ACSF included 10 μM NBQX and 3 μM CGP-55845 to block AMPA-type glutamate receptors and GABA_B_ receptors, respectively. For a subset of experiments (see text), the ACSF included (in μM): 50 picrotoxin, 10 or 0.02 gabazine, 1 TTA-A2, 10 RO4938581, 1 Compound A, or 2 DHPG. TTA-A2 was a gift from Bruce Bean, Harvard Medical School. RO4938581 and Compound A were synthesized and contributed by Hoffman-La Roche (Basel, Switzerland). All other compounds were purchased from Tocris.

### Radioligand binding and electrophysiological assays of recombinant receptors

The cDNAs encoding different rat and human GABA_A_R subunits (α1, α2, α3, α5, β2, β3 and γ2) were subcloned into the polylinker of the pcDNA3.1 vector (Invitrogene, USA) by standard techniques for transiently transfecting HEK293-F cells (ThermoFisher). Cells were transiently transfected with the plasmids containing the desired GABA_A_R subunit cDNAs (α, β, γ at a 1:1:1 ratio) using retro-inverso dioleoylmelittin (riDOM) 0.2 mg/nl (synthesized at Hoffman-La Roche) and PEI 25 kD 0.67 mg/ml. At 48 h post-transfection, the cells were harvested for membrane preparation and radioligand binding assays as described previously (Ballard et al., 2009). Briefly, the inhibition of 1 nM [^3^H]-flumazenil binding by Compound A was measured in membranes expressing either α1β3γ2, α2β3γ2, α3β3γ2 or α5β3γ2 receptors. Non-specific binding was determined in the presence of 10 μM diazepam. Affinity values were calculated using Excel-Fit (Microsoft).

For electrophysiology studies, HEK293 cells stably expressing α1β2γ2, α2β3γ2, α3β3γ2 or α5β3γ2 receptor subtypes were obtained and maintained as previously described (Ballard et al., 2009). Cells were plated on glass coverslips, transferred to a chamber on the stage of a Nikon Diaphot 300 inverted microscope, and continuously superfused with a solution consisting of (in mM) 150 NaCl, KCl, 1.2 CaCl_2_, 1 MgCl_2_, 10 HEPES and 30 sucrose, adjusted to pH 7.4. Glass electrodes (2-3 MΩ) were filled with a solution containing (in mM) 140 CsCl, 10 HEPES, 11 EGTA, 1 CaCl_2_, 1 MgCl_2_, 4 Mg-ATP and 25 sucrose, pH adjusted to 7.2 with CsOH. Whole-cell voltage-clamp recordings were performed at −60 mV using a MultiClamp 700A amplifier (Axon Instruments). GABA in the presence or absence of drugs was applied to the cell for 1 s in 1-minute intervals using a multi-barreled micro-applicator (RSC-200, Biologic Science Instruments). For each experiment, at least 3 GABA control applications were generated, and only cells showing stable responses were selected for drug testing. Maximally effective concentrations of midazolam were applied for each GABA_A_R subtype as a positive control. Concentration-response curves were generated by applying increasing concentrations of each test drug to the same cell until a maximum response was observed (usually 2-3 times).

### Computational Modeling

Single neurons were modeled in the Neuron environment (Hines and Carnevale, 1997) as a 20 μm radius soma attached to a 300 μm dendrite evenly divided into 17 compartments with 0.5 μm diameter. Voltage-activated sodium, potassium, and calcium channels were taken from a previous model of a cortical pyramidal cell (Kampa and Stuart, 2006). A leak conductance with a reversal potential of −60 mV and a tonic GABAergic conductance with a reversal potential of – 70 mV were also included. All conductances were expressed uniformly in the cell membrane. Passive membrane properties and conductance values are specified in Supplemental Figure 2, and the entire model is available on ModelDB (accession 258630). To evaluate GABAergic modulation of AP-evoked calcium signals, a 2 ms 2 nA current was injected at the soma to evoke an action potential that propagated throughout the dendrite. ΔCa^2+^ was defined as the baseline-subtracted, integrated current through the included calcium channels from 1-20 ms following the AP. Simulations were run in the presence and absence of the tonic GABAergic conductance.

## Acknowledgements

The authors wish to thank members of the Higley laboratory and Dr. Jessica A. Cardin for helpful comments during the preparation of this manuscript, the Yale Center for Research Computing for support with the Yale Farnam high performance cluster, Henner Knust for compound synthesis, Chiristian Miscenic and Marcello Foggetta for cell transfections and membrane preparations, Judith Lengyel, Gregoire Friz, and Maria Karg for cell line generation and radioligand binding assays, and Marie Claire Pflimlin for support with electrophysiological characterization of Compound A selectivity.

## Authors contributions

Experiments were conceived and designed by CQC and MJH. Experimental data were acquired by CQC and FN. Modeling data were generated by MJ and TMM. Analyses were carried out by CQC, TMM, FN, and MJH. Work was directed and supervised by FK, MCH, and MJH. Manuscript was written by CQC, TMM, MCH, and MJH.

## Data and materials availability

All data are presented in the paper and supplementary materials. The datasets generated during the current study are available from the corresponding author on reasonable request. The computational model is available through the ModelDB repository (http://modeldb.yale.edu/258630).

## Supplementary Materials

## Supplemental Figure Legends

**Supplemental Figure 1.**
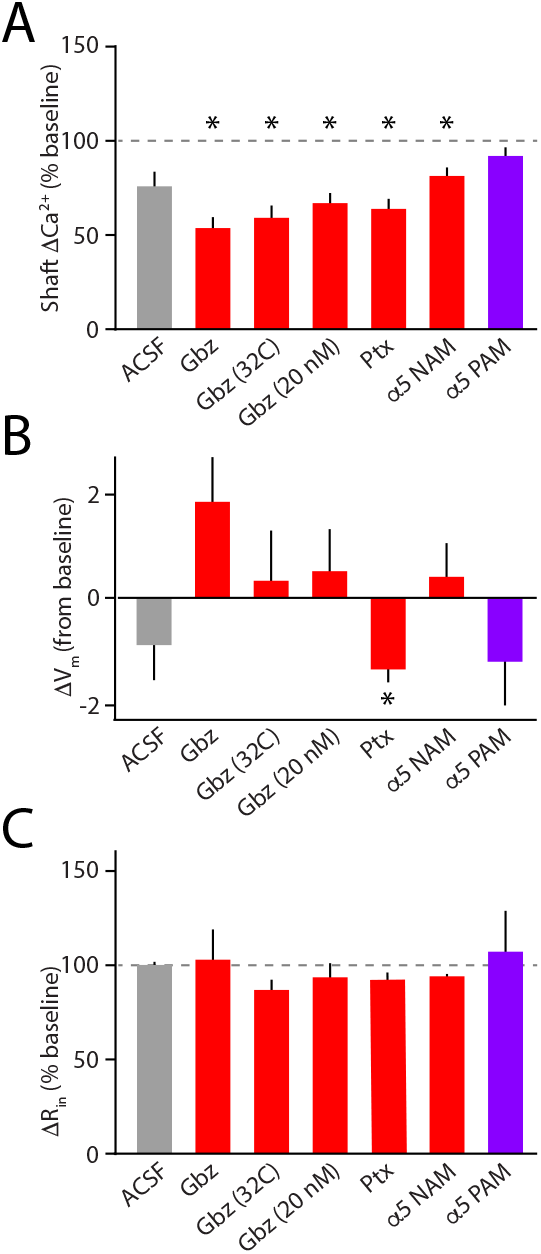
GABAergic modulation of dendritic calcium signaling occurs without alteration in somatic passive membrane properties. **A**, Population data indicating the change in dendritic shaft ΔCa^2+^ following flow-in of ACSF, 10 μM gabazine, 10 μM gabazine at 32°C, 20 nM gabazine, 10 μM picrotoxin, 10 μM RO4938581 (NAM), or 1 μM Compound A (PAM). Error bars indicate SEM. * indicates p<0.05 relative to baseline (dashed line). **B**, As in (A), indicating change in somatic membrane potential. **C**, As in (A), indicating change in somatic input resistance.

**Supplemental Figure 2.**
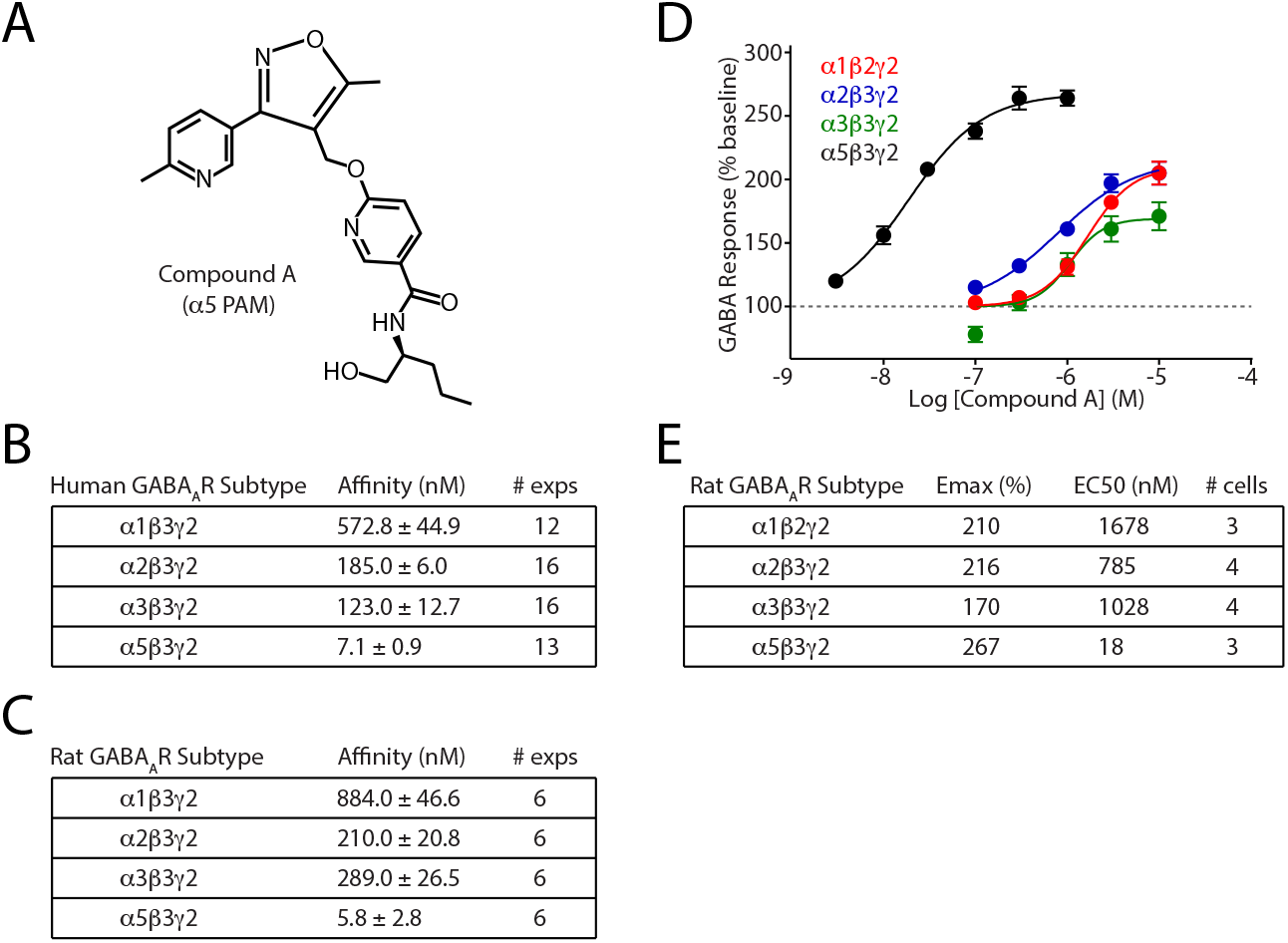
Pharmacology of the selective α5 positive allosteric modulator Compound A. **A**, Schematic illustrating the molecular structure of Compound A, the selective α5 positive allosteric modulator. **B**, Table indicating the affinity of Compound A for different human GABA_A_R subtypes determined by inhibition of [^3^H]-flumazenil binding. **C**, Table indicating the affinity of Compound A for different rat GABA_A_R subtypes determined by inhibition of [^3^H]-flumazenil binding. **D**, Population data showing response curves (mean ± SEM) for GABA-evoked whole-cell currents through different heterologous rat GABA_A_Rs in the presence of varying concentrations of Compound A. Data were fitted with a sigmoid function. **E**, Table indicating the curve parameters derived from data in (E).

**Supplemental Figure 3.**
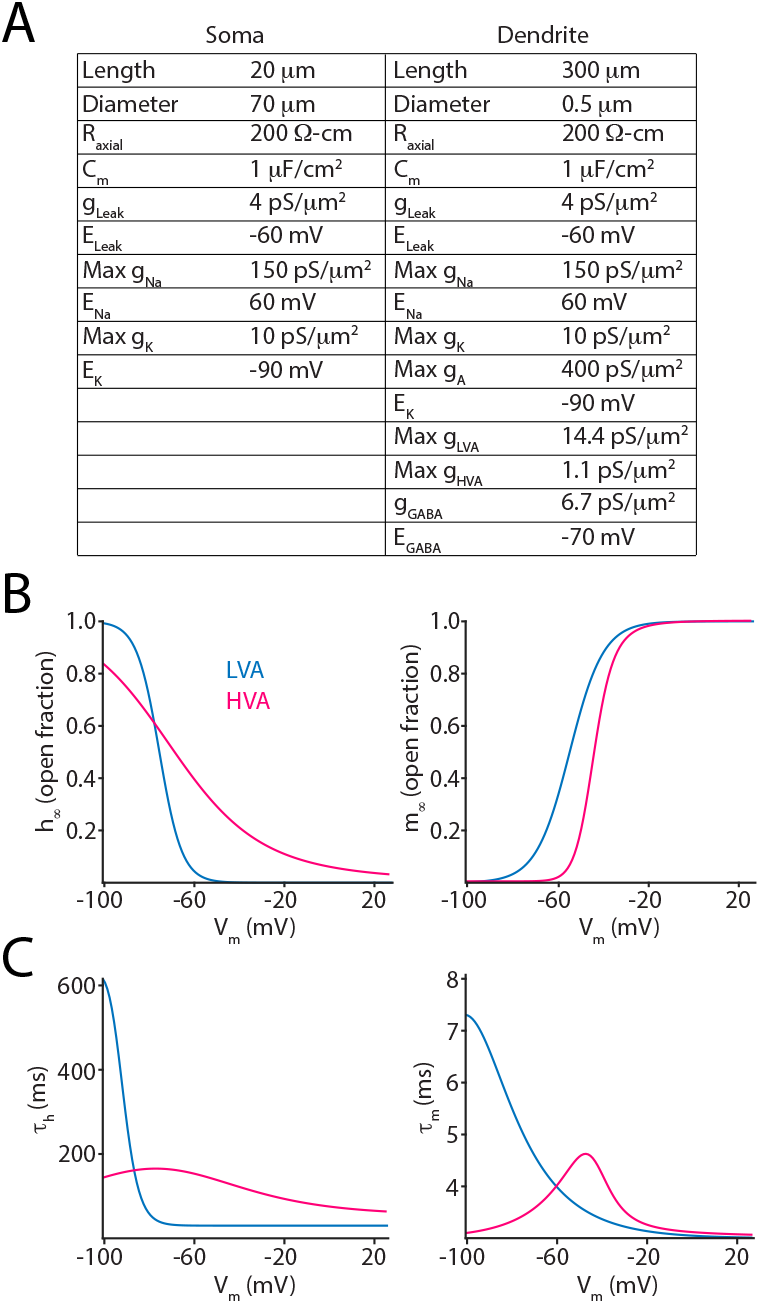
Computational model parameters for investigating the consequences of tonic inhibition on dendritic calcium signaling. **A**, Table indicating the passive membrane properties and channel conductances for the somatic and dendritic compartments of the model. **B**, Relationship between membrane voltage and steady-state inactivation (left) or activation (right) for the LVA (blue) and HVA (pink) channels present in the model. **C,** Relationship between membrane potential and gating kinetics for inactivation (left) and activation (right) gates for the LVA (blue) and HVA (pink) channels present in the model.

## References

Bai, D., Zhu, G., Pennefather, P., Jackson, M.F., MacDonald, J.F., and Orser, B.A. (2001). Distinct functional and pharmacological properties of tonic and quantal inhibitory postsynaptic currents mediated by gamma-aminobutyric acid(A) receptors in hippocampal neurons. Mol Pharmacol 59, 814–824.

Ballard, T.M., Knoflach, F., Prinssen, E., Borroni, E., Vivian, J.A., Basile, J., Gasser, R., Moreau, J.L., Wettstein, J.G., Buettelmann, B., et al. (2009). RO4938581, a novel cognitive enhancer acting at GABAA alpha5 subunit-containing receptors. Psychopharmacology (Berl) 202, 207–223.

Braudeau, J., Delatour, B., Duchon, A., Pereira, P.L., Dauphinot, L., de Chaumont, F., Olivo-Marin, J.C., Dodd, R.H., Herault, Y., and Potier, M.C. (2011). Specific targeting of the GABA-A receptor alpha5 subtype by a selective inverse agonist restores cognitive deficits in Down syndrome mice. J Psychopharmacol 25, 1030–1042.

Caraiscos, V.B., Elliott, E.M., You-Ten, K.E., Cheng, V.Y., Belelli, D., Newell, J.G., Jackson, M.F., Lambert, J.J., Rosahl, T.W., Wafford, K.A., et al. (2004). Tonic inhibition in mouse hippocampal CA1 pyramidal neurons is mediated by alpha5 subunit-containing gamma-aminobutyric acid type A receptors. Proc Natl Acad Sci U S A 101, 3662–3667.

Cardin, J.A. (2018). Inhibitory Interneurons Regulate Temporal Precision and Correlations in Cortical Circuits. Trends Neurosci 41, 689–700.

Carreno, F.R., Lodge, D.J., and Frazer, A. (2020). Ketamine: Leading us into the future for development of antidepressants. Behav Brain Res 383, 112532.

Chalifoux, J.R., and Carter, A.G. (2011). GABAB receptor modulation of voltage-sensitive calcium channels in spines and dendrites. J Neurosci 31, 4221–4232.

Chevaleyre, V., Takahashi, K.A., and Castillo, P.E. (2006). Endocannabinoid-mediated synaptic plasticity in the CNS. Annu Rev Neurosci 29, 37–76.

Chiu, C.Q., Barberis, A., and Higley, M.J. (2019). Preserving the balance: diverse forms of long-term GABAergic synaptic plasticity. Nat Rev Neurosci 20, 272–281.

Chiu, C.Q., Lur, G., Morse, T.M., Carnevale, N.T., Ellis-Davies, G.C., and Higley, M.J. (2013). Compartmentalization of GABAergic inhibition by dendritic spines. Science 340, 759–762.

Chiu, C.Q., Martenson, J.S., Yamazaki, M., Natsume, R., Sakimura, K., Tomita, S., Tavalin, S.J., and Higley, M.J. (2018). Input-Specific NMDAR-Dependent Potentiation of Dendritic GABAergic Inhibition. Neuron 97, 368–377 e363.

Cichon, J., and Gan, W.B. (2015). Branch-specific dendritic Ca(2+) spikes cause persistent synaptic plasticity. Nature 520, 180–185.

Collinson, N., Kuenzi, F.M., Jarolimek, W., Maubach, K.A., Cothliff, R., Sur, C., Smith, A., Otu, F.M., Howell, O., Atack, J.R., et al. (2002). Enhanced learning and memory and altered GABAergic synaptic transmission in mice lacking the alpha 5 subunit of the GABAA receptor. J Neurosci 22, 5572–5580.

Dominguez, S., Fernandez de Sevilla, D., and Buno, W. (2016). Muscarinic Long-Term Enhancement of Tonic and Phasic GABAA Inhibition in Rat CA1 Pyramidal Neurons. Front Cell Neurosci 10, 244.

Donegan, J.J., Boley, A.M., Yamaguchi, J., Toney, G.M., and Lodge, D.J. (2019). Modulation of extrasynaptic GABAA alpha 5 receptors in the ventral hippocampus normalizes physiological and behavioral deficits in a circuit specific manner. Nature communications 10, 2819.

Engin, E., Benham, R.S., and Rudolph, U. (2018). An Emerging Circuit Pharmacology of GABAA Receptors. Trends Pharmacol Sci 39, 710–732.

Farrant, M., and Nusser, Z. (2005). Variations on an inhibitory theme: phasic and tonic activation of GABA(A) receptors. Nat Rev Neurosci 6, 215–229.

Fritschy, J.M., and Mohler, H. (1995). GABAA-receptor heterogeneity in the adult rat brain: differential regional and cellular distribution of seven major subunits. J Comp Neurol 359, 154–194.

Gogolla, N., Leblanc, J.J., Quast, K.B., Sudhof, T., Fagiolini, M., and Hensch, T.K. (2009). Common circuit defect of excitatory-inhibitory balance in mouse models of autism. J Neurodev Disord 1, 172–181.

Groen, M.R., Paulsen, O., Perez-Garci, E., Nevian, T., Wortel, J., Dekker, M.P., Mansvelder, H.D., van Ooyen, A., and Meredith, R.M. (2014). Development of dendritic tonic GABAergic inhibition regulates excitability and plasticity in CA1 pyramidal neurons. J Neurophysiol 112, 287–299.

Gu, X., Zhou, L., and Lu, W. (2016). An NMDA Receptor-Dependent Mechanism Underlies Inhibitory Synapse Development. Cell Rep 14, 471–478.

Hauser, J., Rudolph, U., Keist, R., Mohler, H., Feldon, J., and Yee, B.K. (2005). Hippocampal alpha5 subunit-containing GABAA receptors modulate the expression of prepulse inhibition. Mol Psychiatry 10, 201–207.

Hayama, T., Noguchi, J., Watanabe, S., Takahashi, N., Hayashi-Takagi, A., Ellis-Davies, G.C., Matsuzaki, M., and Kasai, H. (2013). GABA promotes the competitive selection of dendritic spines by controlling local Ca2+ signaling. Nat Neurosci 16, 1409–1416.

Higley, M.J. (2014). Localized GABAergic inhibition of dendritic Ca(2+) signalling. Nat Rev Neurosci 15, 567–572.

Hines, M.L., and Carnevale, N.T. (1997). The NEURON simulation environment. Neural computation 9, 1179–1209.

Jacob, T.C. (2019). Neurobiology and Therapeutic Potential of alpha5-GABA Type A Receptors. Front Mol Neurosci 12, 179.

Jiang, L., Kang, D., and Kang, J. (2015). Potentiation of tonic GABAergic inhibition by activation of postsynaptic kainate receptors. Neuroscience 298, 448–454.

Kampa, B.M., and Stuart, G.J. (2006). Calcium spikes in basal dendrites of layer 5 pyramidal neurons during action potential bursts. J Neurosci 26, 7424–7432.

Lee, V., and Maguire, J. (2014). The impact of tonic GABAA receptor-mediated inhibition on neuronal excitability varies across brain region and cell type. Front Neural Circuits 8, 3.

Magnin, E., Francavilla, R., Amalyan, S., Gervais, E., David, L.S., Luo, X., and Topolnik, L. (2019). Input-Specific Synaptic Location and Function of the alpha5 GABAA Receptor Subunit in the Mouse CA1 Hippocampal Neurons. J Neurosci 39, 788–801.

Marlin, J.J., and Carter, A.G. (2014). GABA-A receptor inhibition of local calcium signaling in spines and dendrites. J Neurosci 34, 15898–15911.

Paine, T.A., Chang, S., and Poyle, R. (2020). Contribution of GABAA receptor subunits to attention and social behavior. Behav Brain Res 378, 112261.

Pologruto, T.A., Sabatini, B.L., and Svoboda, K. (2003). ScanImage: flexible software for operating laser scanning microscopes. Biomed Eng Online 2, 13.

Prevot, T.D., Li, G., Cook, J.M., and Sibille, E. (2019). Insight into Novel Treatment for Cognitive Dysfunctions across Disorders. ACS Chem Neurosci 10, 2088–2090.

Schulz, J.M., Knoflach, F., Hernandez, M.C., and Bischofberger, J. (2018). Dendrite-targeting interneurons control synaptic NMDA-receptor activation via nonlinear alpha5-GABAA receptors. Nature communications 9, 3576.

Scimemi, A., Semyanov, A., Sperk, G., Kullmann, D.M., and Walker, M.C. (2005). Multiple and plastic receptors mediate tonic GABAA receptor currents in the hippocampus. J Neurosci 25, 10016–10024.

Stell, B.M., and Mody, I. (2002). Receptors with different affinities mediate phasic and tonic GABA(A) conductances in hippocampal neurons. J Neurosci 22, RC223.

